# Genes and pathways determining flowering time variation in temperate adapted sorghum

**DOI:** 10.1101/2024.12.12.628249

**Authors:** Harshita Mangal, Kyle Linders, Jonathan Turkus, Nikee Shrestha, Blake Long, Ernst Cebert, Xianyan Kuang, J. Vladimir Torres-Rodriguez, James C. Schnable

## Abstract

The timing of the transition from vegetative to reproductive growth is determined by a complex genetic architecture integrating signals from a diverse set of external and internal stimuli and plays a key role in determining plant fitness and adaptation. However, significant divergence in the identities and functions of many flowering time pathway components has been reported among plant species. Here we employ a combination of genome and transcriptome wide association studies to identify genetic determinants of variation in flowering time across multiple environments in a large panel of primarily photoperiod-insensitive sorghum (*Sorghum bicolor*), a major crop that has, to date, been the subject of substantially less genetic investigation than its relatives. Gene families that form core components of the flowering time pathway in other species, FT-like and SOC1-like genes, appear to play similar roles in sorghum, but the genes identified are not orthologous to the primary FT-like or SOC1-like genes which play similar roles in related species. The aging pathway appears to play a significant role in determining non-photoperiod determined variation in flowering time in sorghum. Two components of this pathway were identified in a transcriptome wide association study, while a third was identified via genome wide association. Our results demonstrate that, while the functions of larger gene families are conserved, functional data from even closely related species is not a reliable guide to which gene copies will play roles in determining natural variation in flowering time.

## Introduction

Flowering time is a complex phenotype that is determined by the integration of a large number of distinct signaling pathways and plays a critical role in determining fitness and environmental adaption in both crops and wild species. In the two plant systems where the study of genes involved in flowering time is most advanced – rice and arabidopsis – a number of commonalities are observed, particularly the key role of different members of the phosphatidyl ethanolamine-binding proteins gene family – *FT* in arabidopsis (*Arabidopsis thaliana*) and *Hd3a* in rice (*Oryza sativa*) (Kobayashi et al., 1999). One of the primary flowering time pathways depends on sensing of day length (photoperiod). Photoperiod sensing will allow plants which germinate at different times to consistently target reproduction to times of the year with optimal conditions. However, in domesticated crops, photoperiod sensitivity typically represented a barrier to the dispersal of domesticated crops into new regions, while the loss of seed dormancy and controlled planting dates also associated with domestication largely eliminate the need for photoperiod sensing. As a result, the loss or attenuation of the role of the photoperiod in determining flowering time has been a common pattern among globally distributed crops including maize (*Zea mays*), rice, soybean (*Glycine max*), and tomato (*Solanum lycopersicum*) (Huang et al., 2018, Zong et al., 2021, Lin et al., 2021, Song et al., 2020). Beyond photoperiod a diverse set of other environment sensing or time dependent genetic pathways have been showed to influence flowering time in the model plant arabidopsis including the shade avoidance/light quality pathway, vernalization pathway, autonomous pathway, temperature-sensing pathway, aging pathway, and GA pathway (Freytes et al., 2021, Cao et al., 2021).

Sorghum, (*Sorghum bicolor*) was originally domesticated in the tropical latitudes of Africa and tropical adapted sorghum lines are typically photoperiod sensitive, with flowering induced by shorter day lengths (Quinby, 1967). In temperate latitudes many tropical sorghum varieties will fail to flower early enough to complete their life cycle prior to being killed by frost. Temperate adapted sorghum typically carries loss of function alleles for one or more of several well-characterized maturity genes which act at different points in the photoperiod sensing flowering time pathway (Quinby, 1967). *Maturity1* (*Ma1*) is associated with the largest impact on sorghum flowering under non-inductive long day conditions (Quinby, 1967) and corresponds to *SbPRR37* (Sobic.006G057866) (Murphy et al., 2011) a homolog of the green revolution gene *Ppd1* which converted photoperiod sensitive wheat to photoperiod insensitivity (Beales et al., 2007). At least seven independent loss of function alleles of *Ma1* have been observed in photoperiod insensitive sorghum suggesting that selection for flowering under long-day conditions repeatedly selected for loss of function of this gene (Murphy et al., 2011, Klein et al., 2015). The photoperiod insensitive allele of *Maturity2* is associated with greater expression of the *FT* homolog *SbFT8* under long-day conditions and is believed to correspond to Sobic.002G302700 which encodes a SMYD domain containing protein (Casto et al., 2019). *Maturity3* and *Maturity5* are thought to correspond to Sobic.001G394400 and Sobic.001G087100, which encode the red/far-red light sensing proteins phytochromeB and phytochromeC respectively (Childs et al., 1997, Yang et al., 2014). *Maturity6* has been mapped to Sobic.006G004400 which encodes a CCT domain containing protein orthologous to the rice flowering time gene *GHD7* and the maize flowering time genes *ZmCCT9* and *ZmCCT10* (Murphy et al., 2014, Huang et al., 2018) and not corresponding to any annotated gene in arabidopsis (Xue et al., 2008). In addition to functional characterization of multiple loss of function alleles for some classical sorghum maturity genes (Murphy et al., 2011, Klein et al., 2015), analysis of resequencing data from large sorghum populations suggests additional loss or modification of function alleles may be present for a number of these loci (Grant et al., 2023). The presence of multiple independent loss of function alleles for the same gene can present challenges for genome-wide association studies as no one genetic marker is likely to separate functional and non-functional haplotypes. However, the flowering times of sorghum genotypes carrying the same alleles of known maturity genes can vary substantially, indicating many other uncharacterized loci also play a role in determining flowering time in sorghum.

The characterized sorghum maturity genes largely function via the photoperiod sensitive flowering time pathway which distinguishes tropical and temperate adapted sorghum varieties. Efforts to increase the genetic diversity accessible to temperate sorghum breeders motivated the creation of large numbers of sorghum conversion lines where a single temperate adapted donor line was employed to introgress genes for short stature and photoperiod insensitivity into hundreds of tropically adapted sorghum lines from around the globe (Stephens et al., 1967). However, even after selecting for photoperiod insensitivity, sorghum conversion lines exhibit a wide range of flowering times. The members of the sorghum association panel, a diversity panel composed primarily of sorghum conversion lines (Casa et al., 2008), flowered between 48 and 87 days after planting when grown in Lincoln, Nebraska (Grzybowski et al., 2022) and similar wide ranges of maturity times were reported for field trials of the sorghum association panel in other locations (Zhao et al., 2016, Wei et al., 2024). While the sorghum association panel was been employed extensively for genome wide association studies of a range of traits, only two studies had reported significant genome wide association hits for flowering time in the sorghum association panel prior to the cutoff for a 2021 metaanalysis (Mural et al., 2021), one of which identified five signals on the edge of statistical significance on sorghum chromosomes 5, 8, 9, and 10, none of which were associated with characterized sorghum flowering time genes (Zhao et al., 2016) and a second study identified a signal in the vicinity of *Ma1* as well two additional signals on sorghum chromosomes 1 and 10 which did not correspond to known flowering time genes (Miao et al., 2020). A joint analysis of flowering time variation in the sorghum association panel across 14 locations in the USA identified three loci associated with plasticity in flowering time, including one in the region of *Ma1* and two signals on chromosomes 1 and 5, but no signals significantly associated with non-environmentally plastic variation in flowering time (Wei et al., 2024).

The lack of progress in identifying and characterizing the regulators of variation in flowering time in temperate adapted sorghum is likely not attributable to the difficulty in accurately scoring this trait as measurements of flowering time in sorghum tend to be highly heritable (Zhao et al., 2016, Mural et al., 2021, Wei et al., 2024). We speculated that one or both of two factors may explain this lack of progress. Firstly, the size and structure of the sorghum association panel, which consists of approximately 400 unique genotypes of which 300-350 are employed in most studies (Casa et al., 2008, Mural et al., 2021) may simply not be large enough to link genomic intervals to variation in flowering time with sufficient statistical significance. Second, genome wide association studies may not properly capture variation attributable to three or more distinct haplotypes at the same locus, a situation likely to exist if the determinants of non-photoperiod sensitive flowering time variation include genes where multiple independent loss of function alleles have been selected (Torres-Rodríguez et al., 2025), as appears to be the case for at least some of the sorghum maturity genes. Here we employ a larger set of 738 temperate adapted and largely photoperiod insensitive sorghum conversion lines drawn from both the sorghum association panel (Casa et al., 2008) and the sorghum diversity panel (Griebel et al., 2021) to identify genomic intervals and candidate genes associated with variation in flowering time in temperate adapted sorghum under long day conditions. A combination of genome wide association studies and transcriptome wide association studies implicate the aging flowering time pathway, originally characterized in maize and arabidopsis as playing a key role in determining flowering time variation in temperate adapted sorghum across multiple environments.

## Methods

### Field Experiments

A set of 915 sorghum genotypes drawn from both the Sorghum Association Panel (Casa et al., 2008) and the Sorghum Diversity Panel (Griebel et al., 2021) was evaluated in a field experiment conducted at the University of Nebraska-Lincoln’s Havelock Research Farm in Lincoln, NE (40.86, −96.60) in 2021. The field design consisted of two replicated blocks, one block consisted of a 968 single-row plot and another 971 single-row plot resulting in 1,939 plots. Each block included a single entry for each of 915 genotypes with the remaining plots consisting of replicates of the check genotypes Tx430 and BTx623. Each plot consisted of a single 60-inch (approximately 1.5 meter) row. Interrow spacing and spacing between sequential plots were both 30 inches (approximately 0.75 meters). Within rows, the sorghum seeds were planted at a target spacing of 3 inches (approximately 7.6 centimeters) for a target plant density of 21 sorghum plants per row. Before planting, nitrogen was applied at a rate of 80 lbs/acre (approximately 90 kilograms per hectare). The planting of all plots was conducted on a single day, May 25th, 2021.

In a second field experiment, a set of 392 sorghum genotypes drawn from the Sorghum Association Panel(Casa et al., 2008) were evaluated at Alabama A&M University’s Winfred Thomas Agriculture Research Station (34.88 and −86.55) in 2022. The set of sorghum genotypes evaluated included 365 shared with the Lincoln, NE experiment. The field design consisted of three replicated blocks with each block containing 432 plots. Each block included a single entry for each of 392 sorghum genotypes with the remaining plots consisting of replicates of the check genotypes Tx430 and BTx623. Each plot consisted of two parallel 60-inch (approximately 1.5 meters) rows separated by 30-inch (approximately 0.75 meters) with 30-inch (approximately 0.75 meters) alleyways between sequential plots. The target planting density was 21 seeds per row (42 seeds per plot). In Alabama poultry litter fertilizer was applied prior to planting and followed with a synthetic nitrogen fertilizer application at a rate of 80 lbs/acre (approximately 90 kilograms per hectare) approximately three weeks after planting (June 26th). Planting was conducted over a three-day period from May 31st to June 2nd, 2022.

Flowering time was scored in both experiments. In Nebraska, flowering time was defined as the first day that visible anthers were present in at least 50% of the living plants present within a given plot. Out of 1,939 plots grown in the Nebraska field trial, flowering time was successfully recorded for 1,888. In Alabama, the percent of plants within each plot exhibiting visible anthers was scored on a weekly basis, and the date at which 50% of plants in a given plot exhibited anthers was interpolated from these weekly records.

### RNA Extraction and Sequencing

Leaf tissue samples were collected over the course of a single day, August 5th, 2021, with sampling beginning at 9:24 am and ending at 11:51 am. Samples were collected from the “4000s” block of the Lincoln field experiment, which was the more western of the two blocks. For each plot, a single representative plant was selected for sampling, avoiding edge plants where possible. Samples were collected from the fourth leaf, counting downwards from the flag leaf, if present, or counting downwards from the uppermost fully expanded leaf if the flag leaf was not yet present or not yet identifiable. Leaf tissue samples were immediately flash-frozen in liquid nitrogen. Flash-frozen leaf samples were first ground to powder. RNA was extracted from frozen and ground tissue samples using MagMax Plant RNA Isolation Kits (ThermoFisher). RNA sequencing libraries were constructed from extracted RNA samples which passed quality control using TruSeq Stranded mRNA Kits (Illumina). The resulting libraries were pooled and sequenced using an Illumina NovaSeq6000 S4 v1.5 with 2 x 150 base pair read lengths and a target read depth of 20 million total fragments sequenced per library. Sequencing was completed for a total of 748 samples.

### RNA-seq Data Processing

RNA sequencing fastq files were trimmed of low-quality regions and adapters using Trimmomatic (v0.33) with the following parameters ILLUMINACLIP:TruSeq3-PE-2.fa:2:30:10 LEADING:3 TRAILING:3 SLIDINGWINDOW:4:15 MINLEN:35 (Bolger et al., 2014). The trimmed reads were aligned to the the BTx623 v5 reference genome (Sbicolor_730_v5.0.hardmasked.fa) (Institute, 2023) using star/2.7.9a (Dobin et al., 2013) with settings: outFilterMismatchNmax 30, outFilterScoreMinOverLread 0.1, outFilterMatchNminOverLread 0.1, seedSearchStartLmax 20. The resulting alignment results were converted to BAM format, sorted, and indexed using samtools/1.19 (Li et al., 2009). BAM files were modified using the Picard (v.3.0;) (Pic, 2019) using first the AddOrReplaceReadGroups command and the following parameters -RGID 1 -RGLB lib1 -RGPL illumina-RGPU unit1 and then the MarkDuplicate command with the following parameters METRICS_FILE metrics.txt VALIDATION_STRINGENCY SILENT -CREATE_INDEX-true. The marked duplicate reads were split using the SplitNCigarReads function in gatk4/4.1(Brouard et al., 2019) and then reordered using the picard function ReorderSAM with the following parameters -VALIDATION_STRINGENCY SILENT -SEQUENCE_DICTIONARY -CREATE_INDEX true. Variant calls were generated using GATK’s HaplotypeCaller tool with the command HaplotypeCaller -ERC BP_RESOLUTION –all-site-pls true –standard-min-confidence-threshold-for-calling 20. InDels were discarded at this stage, leaving only SNP markers in the dataset. SNPs were filtered using gatk4/4.1’s VariantFiltration function and the following criteria “QD < 2.0” “QUAL < 30.0” “SOR > 3.0” “FS > 60.0” “MQ < 40.0” “MQRankSum < −12.5” “ReadPosRankSum < −8.0”. Subsequent to initial quality filtering, genetic markers with 50% missing data across the set of genotyped sorghum samples, minor allele frequencies *≤* 0.01 or greater than 10% heterozygous genotype calls among samples non-missing genotype values were discarded using bcftools v.1.17 (Danecek et al., 2021). This post-filtering SNP set was imputed using Beagle v5.2 with default parameters (Browning et al., 2018). Ten samples were excluded at this stage which included six additional replicates of the check genotype BTx623, two samples collected from border plots, and the samples “PI 533951” and “PI 534063” for which flowering dates were missed in the Nebraska field experiment. Only SNPs with a minor allele frequency of > 0.05 and frequency of heterozygous genotypes calls < 0.1 among the remaining 738 samples were retained for subsequent analyses, resulting in a final set of 170,072 SNPs.

Gene expression was quantified for each sample using Kallisto (v0.46) (Bray et al., 2016) and a reference database constructed using the “primaryTranscriptOnly” transcript file for Sorghum BTx623 v5 reference genome downloaded from phytozome. Quality trimmed fastq files generated by Trimmomatic as described above were used for quantification. Expression was initially quantified for all 32,160 annotated sorghum genes, however, genes with zero expression across all 738 sorghum genomes were omitted resulting in a final set of 28,159 expressed genes. Initial expression data, quantified in units of transcripts per million (TPM), was rescaled between 0 and 2, with the 5% of samples with the lowest expression set to 0 the 5% of samples with the highest expression set to 2 and the remaining 90% of samples scaled between two based on the formula (SampleTPM/(95thPercentileTPM-5thPercentileTPM))*2 (Li et al., 2021).

### Quantitative Genetic Analysis

Both the genome wide association study and the transcriptome wide association study were conducted using BLUEs calculated from raw plot level flowering time data in either Nebraska or Alabama using the 2D spline method implemented in SpATS (1.0-18) (Rodriguez-Alvarez et al., 2018) with genotypes set as a fixed effect, nseg set to 34,14, block included as an additional fixed effect, row, and column included as additional random effects. Reported heritabilities in this study were calculated by SpATS fitting the same model but with genotype as a random effect. In Alabama, data from a small number of plots where the estimated date at which 50% of plants had flowered was >90 days were excluded prior to fitting the SpATS model and calculating BLUEs (Figure S1). The genome-wide association study was conducted using the FarmCPU algorithm as implemented in rMVP (v1.1.1) (Liu et al., 2016, Yin et al., 2021) with the first three principal components calculated from genetic marker data incorporated as covariates. The analysis was repeated 100 times with phenotype values for a different random subset of 10% of sorghum genotypes masked in each iteration. Each SNP marker that exceeded a statistical significance cutoff of 1.87 * 10^−7^ – corresponding to a Bonferonni corrected p-value of 0.05 considering a total of 170,072 SNPs – in at least one iteration was assigned a resampling model inclusion probability value based on the proportion of all 100 iterations in which that marker exceeded the significance cutoff. Transcriptome-wide association studies were performed using the transformed and rescaled gene expression data described above and the compressed mixed linear model(CMLM) as implemented in GAPIT (v3.4)(Lipka et al., 2012, Zhang et al., 2010). The first three principal components of variation calculated from gene expression data were included as covariates in the transcriptome wide association study. The threshold for statistical significance was the p-value that corresponded to an estimated false discovery rate of 5% calculated using the Benjamini-Hochberg method (Benjamini and Yekutieli, 2001).

### microRNA Analyses

At the time of this analysis the most recent comprehensive set of microRNA annotations for the sorghum genome in miRBase were from BTx623 v3 reference genome (Kozomara et al., 2019). The position of overlapping or nearby gene models were initially used to translate between BTx623 v3 and BTx623 v5 coordinates (Goodstein et al., 2012) with blasts of the annotated stem loop precursor sequence used to confirm positions. Target genes for individual microRNAs were determined using the psRNATarget web server (Dai et al., 2018) and the mature miRNA sequence for each microRNA gene provided by miRBase. Genes identified with a mismatch penality of <2.0 were considered targets of the microRNA for the purposes of our study.

## Results

Between 15.0M and 37.6M fragments per sample were sequenced from mRNA isolated mature leaf tissue collected from 738 sorghum genotypes (mean 24.4M fragments; median 23.8M fragments) (Figure S2). The proportion of RNA-seq reads which could be mapped to transcripts from the BTx623 v5 sorghum reference genome ranged from 58.3% - 94.2% with the exception of one poorly aligning sample. A set of 170,072 SNPs with minor allele frequencies of at least 0.05 and no more than 10% heterozygous genotype calls were identified by aligning these reads to the sorghum reference genome. Sorghum genotypes clustered as expected based on annotated subpopulation identity in a principal component analysis conducted using these SNP markers(Figure S3).

Flowering time can exhibit substantial genotype by environment interactions and, as a result, in order to ensure genetic associations for flowering time are generalizable it is essential to test these associations across multiple environments. Flowering time was scored for diverse sorghum lines in two environments, with an estimated heritability of 0.92 in the first field experiment, conducted in Nebraska, and an estimated heritability of 0.96 in the second field experiment, conducted in Alabama. Spatially corrected BLUEs for flowering time estimated for the two different environments were modestly correlated (pearson r = 0.75) (Figure S1). Consistent with previous limited success mapping genes controlling flowering time in the Sorghum Association Panel, no significant associations were identified when using data only from lines in this panel (Figure S4). However, when data from the full set of 738 genotypes was employed, a resampling based genome wide association study conducted using spatially corrected flowering time data identified six SNPs which were significantly associated with variation in flowering time above a Resampling Model Inclusion Probability (RMIP) threshold of 0.2 (Figure 1A). Three of these signals, located on sorghum chromosomes 3, 6, and 9 exhibited statistically significant association with flowering time scored for a subset of 303 of the same sorghum genotypes in Alabama. The other three trait associated markers identified in the genome wide association study were not significantly associated with variation in flowering time in the Alabama field experiment (Chr03:62,260,337, Chr10:442,685 and Chr05:59978242 p = 0.35, 0.15 and 0.09 respectively) (Table 1).

**Figure 1.**
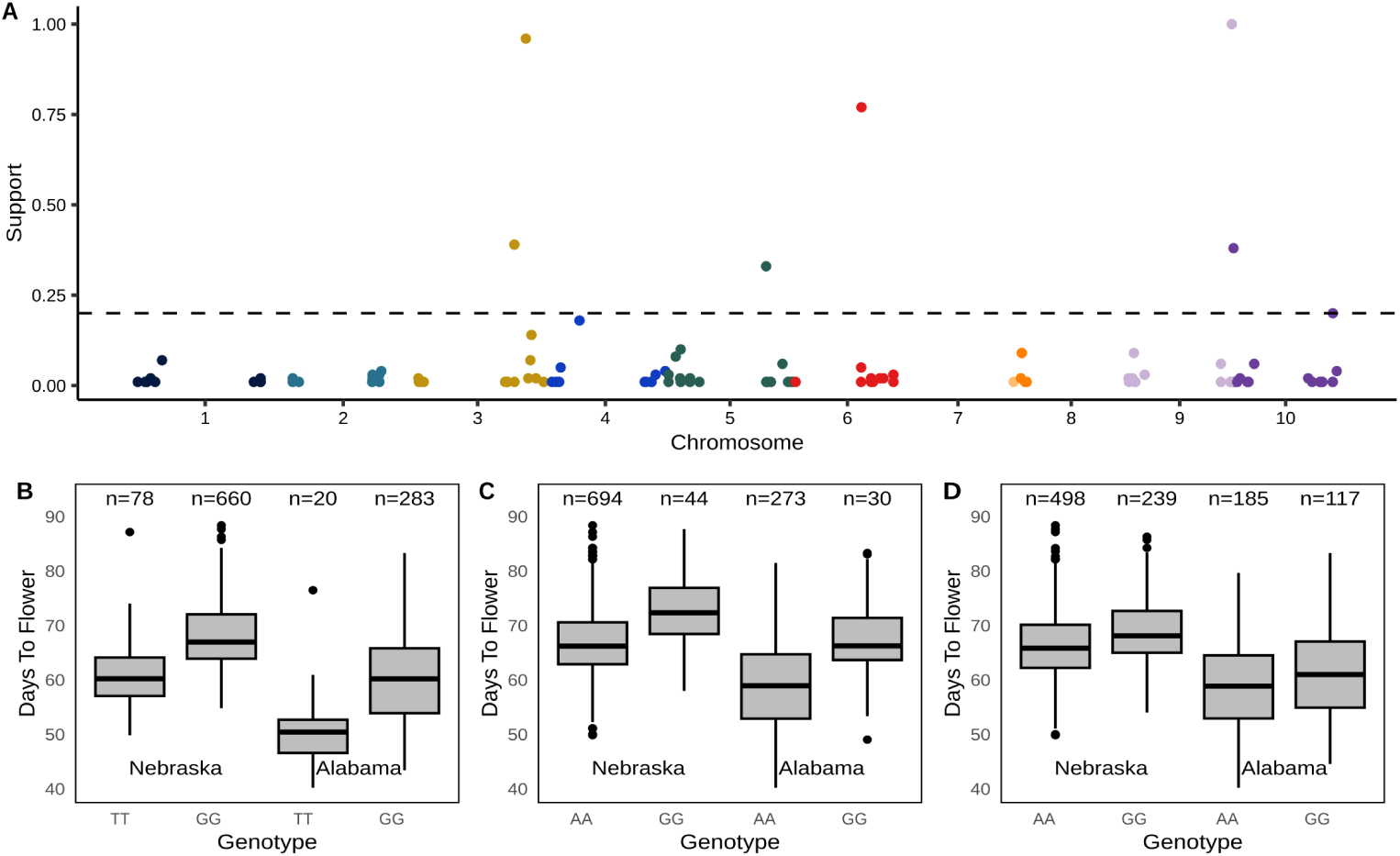
Genetic markers associated with variation in sorghum flowering time across two environments. **A)** Results of a genome wide association study conducted for flowering time measured across 738 sorghum genotypes in Lincoln, Nebraska. Y-axis indicates the resampling model inclusion probability calculated from 100 iterations of the FarmCPU GWAS algorithm. Horizontal dashed line indicates the threshold employed to consider a marker significantly associated with flowering time in this study (RMIP = 0.2). **B)** Difference in best unbiased estimators for flowering time in Nebraska (left) and Alabama (right) between sorghum genotypes homozygous for the T (major) or G (minor) allele at trait associated genetic marker Chr03:69,067,236. Differences were statistically significant in both Nebraska (p = 2.53×10^−9^, MLM-based model) and Alabama (p=5.73×10^−5^, MLM-based model). **C)** Difference in best unbiased estimators for flowering time in Nebraska (left) and Alabama (right) between sorghum genotypes homozygous for the A (major) or G (minor) allele at trait associated genetic marker Chr06:39,694,475.Differences were statistically significant in both Nebraska (p = 3.77×10^−8^, MLM-based model) and Alabama (p=8.20×10^−5^, MLM-based model). **D)** Difference in best unbiased estimators for flowering time in Nebraska (left) and Alabama (right) between sorghum genotypes homozygous for the A (major) or G (minor) allele at trait associated genetic marker Chr09:62,620,720. Differences were statistically significant in both Nebraska (p = 5.93×10^−17^, MLM-based model) and Alabama (p=7.80×10^−5^, MLM-based model).One sorghum variety with a heterozygous genotype call at this locus was excluded from visualization.

**Table 1.**
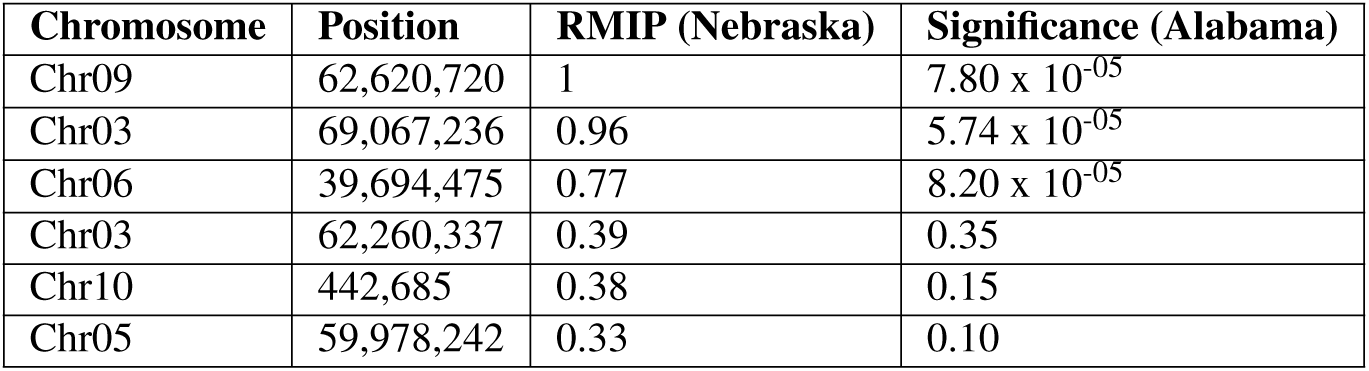
Genetic markers identified via genome wide association study linked to variation in flowering time in sorghum.

| Chromosome | Position | RMIP (Nebraska) | Significance (Alabama) |
| --- | --- | --- | --- |
| Chr09 | 62,620,720 | 1 | $7.80 \times 10^{-05}$ |
| Chr03 | 69,067,236 | 0.96 | $5.74 \times 10^{-05}$ |
| Chr06 | 39,694,475 | 0.77 | $8.20 \times 10^{-05}$ |
| Chr03 | 62,260,337 | 0.39 | 0.35 |
| Chr10 | 442,685 | 0.38 | 0.15 |
| Chr05 | 59,978,242 | 0.33 | 0.10 |

Among the three genetic markers identified in Nebraska and validated in Alabama, Chr03:69,067,236 is located within the sorghum gene model Sobic.003G295300 which encodes a phosphatidylethanolamine-binding protein homologous to the arabidopsis *FT* gene and orthologous the maize gene *zcn12* (Figure 1B, S5A). In this paper we adopt the nomenclature for the phosphatidylethanolamine-binding protein family in sorghum proposed by (Wolabu et al., 2016), who labeled Sobic.003G295300 *SbFT8* (Table S2). The trait associated genetic marker located on sorghum chromosome 6, Chr06:39,694,475 was located in the 5’ UTR of Sobic.006G052100 which encodes a nodulin like protein (Figure 1C, S5B). This gene was not considered a high probability candidate to play a role in determining flowering time and no genes with clear potential links to flowering time were identified within 100 kb of the genetic marker. Chr06:39,694,475 is located approximately two megabases from Sobic.006G057866 *(SbPRR37)*, a gene model thought to correspond to the classical sorghum flowering time gene *Ma1* (Murphy et al., 2011). However, no obvious pattern of elevated linkage disequilibrium was observed between the trait-associated marker and markers adjacent to *SbPRR37* (Figure S5D). The final flowering time associated marker supported by both Nebraska and Alabama datasets was located in a protein coding exon of the gene model Sobic.009G254200, located on sorghum chromosome 9 (Chr09:62,620,720) (Figure 1D, S5C). Sobic.009G254200 encodes a pentatricopeptide repeat domain containing protein with no experimental evidence or obvious mechanistic link to flowering time. Given the elevated linkage disequilibrium exhibited by sorghum generally and observed specifically in the genomic interval surrounding Chr09:62,620,720 specifically we examined the annotated functions of all 25 sorghum gene models located within 100 KB of Chr09:62,620,720, but did not identify any obvious candidates for a role in determining variation in flowering time (Table S1).

A set of 28,159 annotated sorghum genes, out of 32,160 total, exhibited nonzero expression in the expression dataset generated in Nebraska and were employed for transcriptome wide association. In twelve cases, the expression of genes in mature leaf tissue was significantly associated with observed flowering time in Nebraska on a genome-wide basis (FDR <0.05) and for ten of those twelve genes, a significant association was also observed with flowering time in Alabama on a gene-by-gene basis (p <0.05) (Figure 2A; Table 2). A number of genes identified via transcriptome wide association had clear functional links to flowering time, including *SbMADS7* (Sobic.002G010100), which encodes a AP1/FUL/AGL79 MADS-box transcription factor subfamily protein, *SbMADS31* (Sobic.004G003434), which encodes a SOC1 MADS-box transcription factor subfamily protein, and *SbFT2* (Sobic.003G017200), which encodes another member of the phosphatidylethanolamine-binding protein family. However, three of the ten genes whose expression was linked to flowering time variation in sorghum across both environments were annotated as having no known function and did not exhibit significant protein sequence similarity with any annotated genes outside the panicoid grasses. We were able to determine that one of these genes of unknown function without protein sequence similarity to other known genes in fact represented an interval in the sorghum genome encoding a microRNA. Sobic.006G137400 located on sorghum chromosome 6 physically overlapped with the position of the annotation of sbi-MIR156h within miRBase (Kozomara et al., 2019). A second microRNA gene, sbi-MIR156a, overlapped with the position of Sobic.004G065650, another gene model annotated as encoding a protein without homology to known proteins in other species. The association between the expression of Sobic.004G065650 and flowering time approached but did not exceed the threshold for genome wide statistical significance in Nebraska, bit did exceeded the per-gene significance threshold in Alabama. The specific sequence of sbi-MIR156h was determined to target fourteen genes in the sorghum genome encoding SBP transcription factors, including all three SBP genes identified in the TWAS analysis *SbSBP19* (Sobic.002G312300), *SbSBP4* (Sobic.002G312200) and *SbSBP7* (Sobic.004G036900) (Table 3). The expression level of sbi-MIR156h exhibited a negative and non-linear association with the expression of these three transcription factors with spearman correlation values between −0.40 and −0.57 (Figure S6).

**Figure 2.**
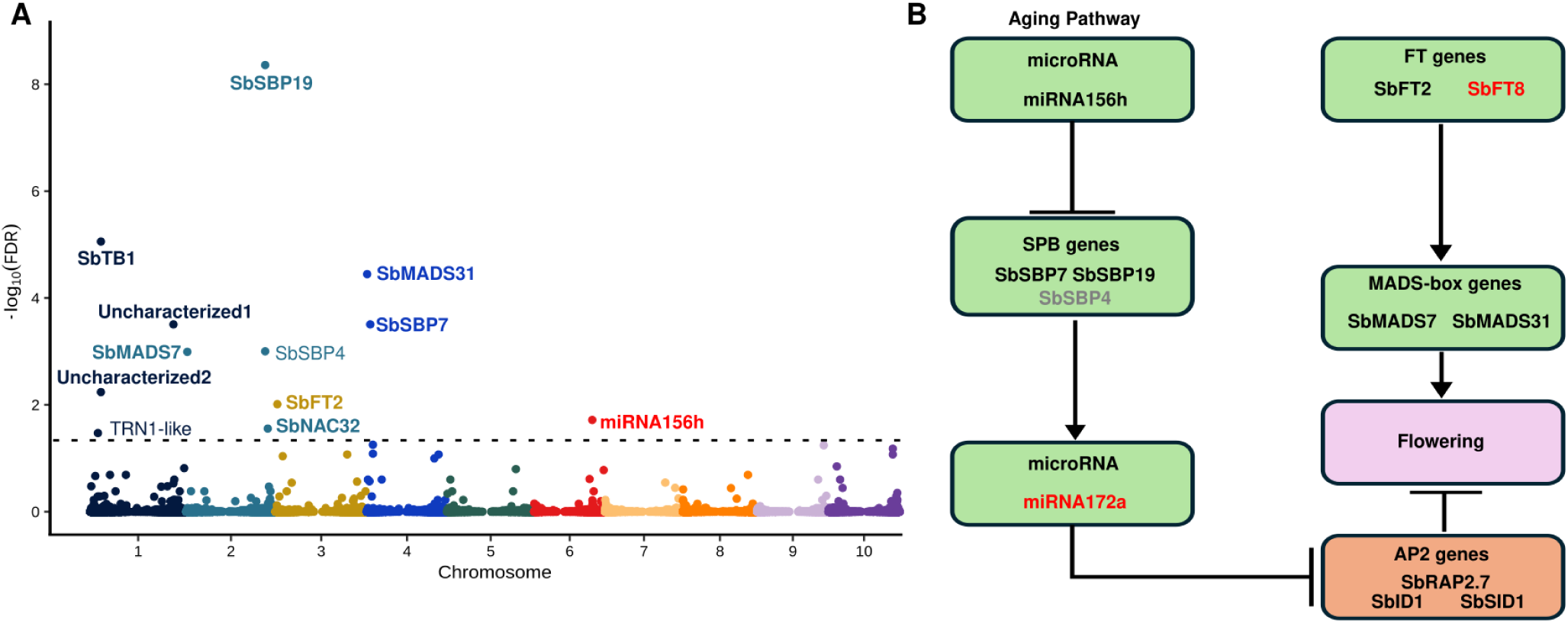
Genes linked to flowering time in sorghum via transcriptome wide association include members of two canonical flowering time pathways. **A)** Results of a transcriptome wide association study conducted for flowering time measured across 738 sorghum genotypes. Each point indicates the significance of the association between the expression of an individual gene and flowering time in Lincoln, Nebraska. Horizontal dashed line indicates the false discovery rate cutoff employed to declare a given association statistically significant (FDR = 0.05). Gene names shown are those assigned in GRASSIUS (transcription factors) (Yilmaz et al., 2009), MirBASE (microRNAs) (Kozomara et al., 2019), or assigned by (Wolabu et al., 2016) (FT genes). Gene names shown in bold were also associated with flowering time in Alabama. **B)** Inferred pathways regulating flowering time in temperate sorghum based on a combination of genome wide association results, transcriptome wide association results, and evidence from the literature. Black names in green ovals were supported by transcriptome associations in both environments. Red names in green ovals were supported by genetic associations in both environments. Grey names in green ovals were supported in only one environment. Ovals shown in orange list computationally confirmed targets of microRNAs identified in our study.

**Table 2.** Sorghum genes linked to flowering time via transcriptome-wide association in descending order of statistical significance.

| GeneID | Symbol | Function | FDR | Significant in Alabama |
| --- | --- | --- | --- | --- |
| Sobic.002G312300 | *SbSBP19* | Transcription factor | $4.37 \times 10^{-9}$ | Yes ($5.38 \times 10^{-5}$) |
| Sobic.001G121600 | *SbTB1* | Transcription factor | $4.37 \times 10^{-9}$ | Yes ($5.023 \times 10^{-3}$) |
| Sobic.004G003434 | *SbMADS31* | Transcription factor | $3.58 \times 10^{-5}$ | Yes ($2.31 \times 10^{-7}$) |
| Sobic.001G414100 | *uncharacterized1* | Unknown | $3.12 \times 10^{-4}$ | Yes ($3.27 \times 10^{-4}$) |
| Sobic.004G036900 | *SbSBP7* | Transcription factor | $3.12 \times 10^{-4}$ | Yes (0.013) |
| Sobic.002G312200 | *SbSBP4* | Transcription factor | $9.95 \times 10^{-4}$ | No (0.20) |
| Sobic.002G010100 | *SbMADS7* | Transcription factor | $1.01 \times 10^{-3}$ | Yes ($1.48 \times 10^{-4}$) |
| Sobic.001G121400 | *uncharacterized2* | Unknown | $5.77 \times 10^{-3}$ | Yes ($1.37 \times 10^{-3}$) |
| Sobic.003G017200 | *SbFT2* | Phosphatidylethanolamine-binding protein | $9.73 \times 10^{-3}$ | Yes ($3.44 \times 10^{-4}$) |
| Sobic.006G137400 | *miRNA156h* | microRNA | 0.019 | Yes ($4.53 \times 10^{-4}$) |
| Sobic.002G342100 | *SbNAC32* | Transcription factor | 0.028 | Yes ($1.04 \times 10^{-3}$) |
| Sobic.001G087800 | *TRN1-like* | Putative signaling protein | 0.033 | No (0.29) |
| Sobic.004G065650 | *miRNA156a* | microRNA | 0.055 | Yes ($9.80 \times 10^{-5}$) |

**Table 3.** Sorghum genes targeted by the two microRNAs linked to flowering time variation in this study.

| microRNA | Target Gene | Gene Description | Mismatch Score |
| --- | --- | --- | --- |
| mir156h | Sobic.004G065700 | sbi-MIR156a | 0 |
| mir156h | Sobic.002G247800 | *SbSBP2* | 1 |
| mir156h | Sobic.002G257900 | *SbSBP3* | 1 |
| mir156h | Sobic.002G312200 | *SbSBP4* | 1 |
| mir156h | Sobic.003G406600 | *SbSBP6* | 1 |
| mir156h | Sobic.004G036900 | *SbSBP7* | 1 |
| mir156h | Sobic.004G058900 | *SbSBP8* | 1 |
| mir156h | Sobic.005G120600 | *SbSBP10* | 1 |
| mir156h | Sobic.006G171000 | *SbSBP11* | 1 |
| mir156h | Sobic.007G193500 | *SbSBP13* | 1 |
| mir156h | Sobic.007G210200 | *SbSBP15* | 1 |
| mir156h | Sobic.010G254200 | *SbSBP18* | 1 |
| mir156h | Sobic.002G312300 | *SbSBP19* | 1 |
| mir156h | Sobic.002G237700 | Inner centromere protein | 1 |
| mir172a | Sobic.001G036800 | ortholog of maize *ids1* | 0.5 |
| mir172a | Sobic.002G083600 | ortholog of maize *sid1* | 0.5 |
| mir172a | Sobic.009G024600 | ortholog of maize *rap2.7* | 0.5 |
| mir172a | Sobic.010G202700 | ortholog of maize *gl15* | 1.5 |

We then searched for microRNAs not represented by annotated gene models in the sorghum BTx623 v5 reference genome. The genomic locus corresponding to sbi-MIR172a is located approximately 40 kilobases upstream from the flowering time associated SNP Chr09:62,620,720 for which no obvious gene candidate has been identified (Figure S7). In sorghum, sbi-MIR172a targets the orthologs of the maize genes *ids1* and *sid1*, both genes involved in floral meristem initiation (Chuck et al., 2008), the ortholog of the maize gene *rap2.7* which acts as repressor of flowering (Salvi et al., 2007), and *glossy15* which promotes juvenile vegetative leaf identity in maize (Lauter et al., 2005) (Table 3). Collectively mir156, SBP transcription factors, and mir172 constitute the major components of the aging flowering time pathway (Figure 2B).

The single gene whose expression was most strongly associated with flowering in our study was *SbSPB19* (Sobic.002G312300). *SbSPB19* and a nearby tandem duplicate, *SbSPB4* (Sobic.002G312200), a gene candidate which was also significantly associated with flowering time, are the respective orthologs of two maize genes, *ZmSPL13* and *ZmSPL29* (Figure 3C) which are targeted by miR156 and whose loss of function alleles result in delayed flowering (Yang et al., 2023). While two sorghum genes encoding PEBP proteins were linked to flowering time – *SbFT8* via GWAS and *SbFT2* via TWAS – neither of these genes were the orthologs of primary functional FT proteins which have been characterized maize (*zcn8*) or rice (*Hd3a*) (Kojima et al., 2002) (Table S2). *SbFT8* is orthologous to the maize gene *zcn12*, while *SbFT2* is orthologous to the maize gene *zcn14* (Figure S8). While *SbFT8* had the second highest average expression of the 13 FT-like genes in the sorghum genome in this study, its expression was not significantly associated with variation in flowering time, even without correction for multiple testing (Table S3). The two MADS box genes whose expression was significantly associated with flowering time variation belonged to different clades within this large gene family. *SbMADS7* (Sobic.002G010100) is the sorghum ortholog of the maize gene *zap1* and encodes a MIKC^C^ type MADS box protein which clusters with the AP1/FUL clade from arabidopsis thought to act as regulators of floral initiation and floral meristem identity. The second MADS box gene significantly associated with variation in sorghum flowering time, *SbMADS31* (Sobic.004G003434) belongs to a clade of MADS box transcription factors in the grasses orthologous to the arabidopsis genes *soc1*, *agl19*, and *agl14*. Three lineages appear to have been present in the common ancestor of rice and sorghum. One lineage contains the maize genes *mads1* and *zag6*, both previously linked to flowering time in maize, and *OsMADS50* (LOC_Os03g03070) which has been shown to function as a floral activator in rice (Lee et al., 2004). A second lineage contained the maize gene *mads76* and the rice gene *OsMADS56* (LOC_Os10g39130) which appears to function as a floral repressor in rice (RYU et al., 2009). The third lineage, which included *SbMADS31*, also included LOC_Os02g01355, conserved as a syntenic location with equivalent intron/exon architecture in rice, however, this conserved gene is apparently completely absent from the maize genome based on both BLAST comparison to annotated maize genes and more detailed examination of the two genomes of the maize genome co-orthologous to the position of Sobic.004G003434 (Figure 3A,B)

**Figure 3.**
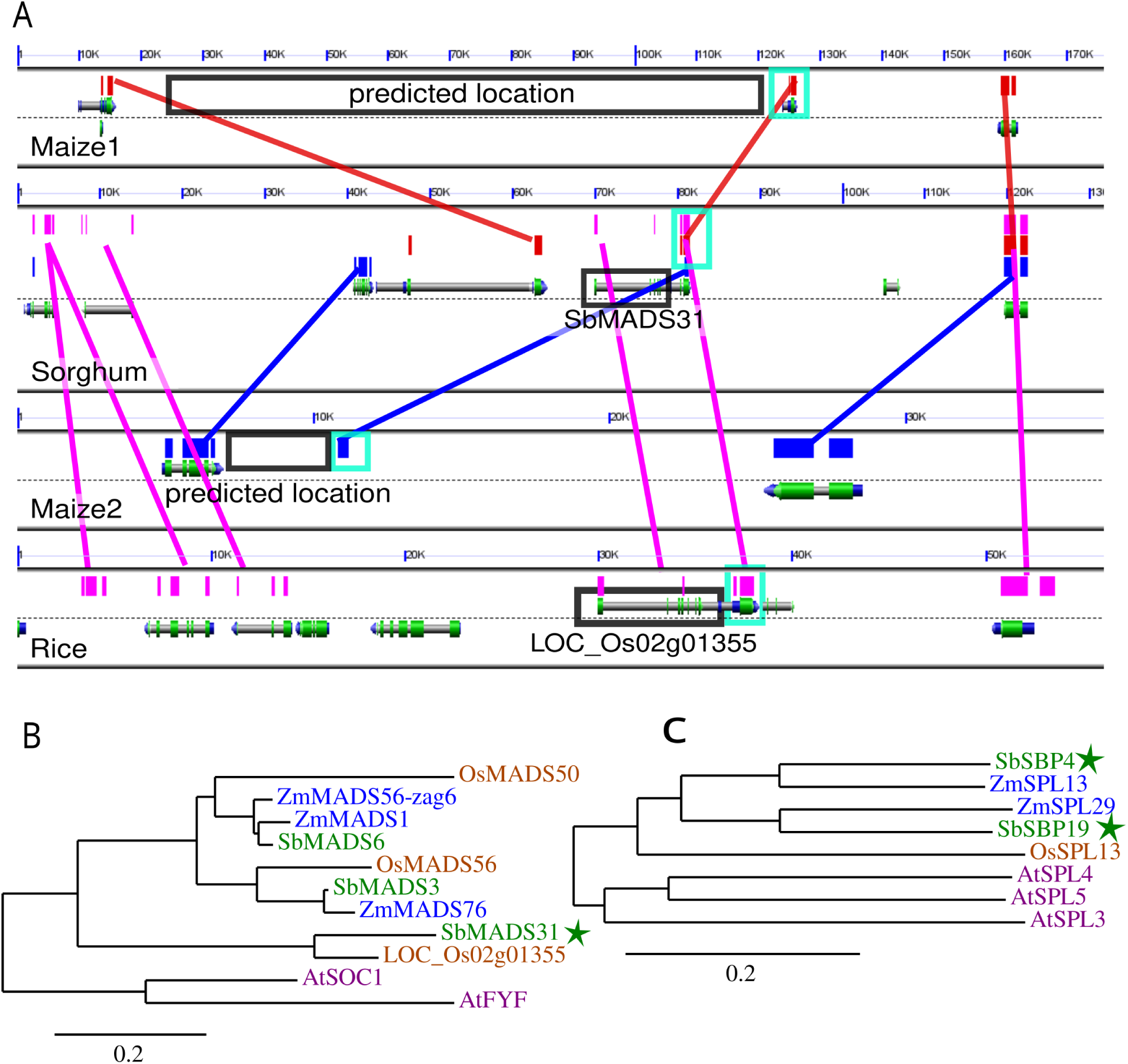
Relation between candidate genes identified in sorghum and characterized genes in other species. **A)** GEvo graphic comparing the genomic interval containing SbMADS31 to the syntenic orthologous genomic interval in rice and the two co-orthologous syntenic regions in the maize genome (Lyons et al., 2008). Red boxes and lines indicate the positions of similar sequences between the maize1 genome and sorghum detected via blastn, blue between sorghum and maize2, and purple between sorghum and rice. Block boxes indicate the position of SbMADS31, its syntenic ortholog in the rice genome, and the two genomic intervals in the maize genome in which orthologs of this sorghum gene would be predicted to be found. **B)** Phylogenetic relationships among the set of SOC1-like MADS box genes found in the sorghum, maize, rice, and arabidopsis genomes. Star indicates the sorghum gene copy linked to variation in flowering time via TWAS in our study. Phylogeny constructed using PhyML using a multiple sequence generated by aligning the nucleotide sequence of the primary transcript for each gene with MUSCLE. **C)** Phylogenetic relationships among the closest relatives of SbSBP19 and SbSBP4 in the maize rice and arabidopsis genomes. Stars indicate the sorghum genes linked to variation in flowering time via TWAS in our study. Phylogeny constructed as described in panel B.

## Discussion

Prior efforts to understand the genetic basis of flowering time variation in sorghum have focused to a significant extent on the six classical maturity genes which appear to play roles in the photoperiod flowering time pathway. When we examined flowering time variation in a photoperiod insensitive sorghum panel under long days none of the loci we identified corresponded to the classical maturity genes, with the potential exception of a signal several megabases removed from *SbPRR37* (Figure S5D), the gene model believed to correspond to *Ma1*(Murphy et al., 2011). Several previous studies conducted with different marker sets at different or only partially overlapping populations of sorghum genotypes have also reported GWAS associations for flowering time in the general vicinity of *Ma1* while farther from *SbPRR37* than would be expected (Higgins et al., 2014, Zhao et al., 2016, Miao et al., 2020, Yang et al., 2014) even given the elevated linkage disequilibrium present in sorghum. This may indicate that the signal reflects functional variation in a gene other than the putative *Ma1* gene model *PRR37* or the difficulty in precisely localizing *Ma1* via GWAS may reflect the large region selected around the *Ma1*/*Dw2* locus as part of the sorghum conversion process (Thurber et al., 2013).

Instead, we identified both a genome wide association signal from *SbFT8*, the sorghum ortholog of *zcn12*, and a transcriptome wide association signal from *SbFT2* the sorghum ortholog of *zcn14*. Of the two sorghum genes identified *SbFT8* is more closely related to the primary FT-like gene linked to flowering time variation in maize(Meng et al., 2011, Wolabu et al., 2016) (Figure S8) but it is not the direct ortholog of that maize gene. *SbFT8* has previously been reported to be expressed at high levels in photoperiod insensitive sorghum leaves but not in photoperiod sensitive sorghum under long day – non-flowering inducing – conditions (Wolabu et al., 2016). The maize ortholog of *SbFT8*, *zcn12*, is competent to promote early flowering when overexpressed in arabidopsis, and variation in the expression of *zcn12* has been linked to flowering time variation in temperate – largely photoperiod insensitive – maize populations phenotyped in both North America and Europe (Torres-Rodríguez et al., 2024, Castelletti et al., 2020). *SbFT2* had not been considered a previous high-priority candidate for the functional ortholog of *FT* in sorghum (Wolabu et al., 2016). However, expression of the maize ortholog of *SbFT2*, *zcn14*, has also been linked to variation in flowering time via transcriptome wide associations study in maize(Torres-Rodríguez et al., 2024) and expression data from tropical photoperiod sensitive maize suggests that this gene may play a role in inducing flowering time in these maize lines(Meng et al., 2011). Our analysis also identified two MADS-box genes associated with variation in flowering time in sorghum. *SbMADS7* is most similar to the MADS box genes *ful*, *ap1* and *cal1* in arabidopsis which function in specifying inflorescence meristem identity, downstream of *soc1*, and the ortholog of the rice gene LOC_Os07g01820 (*MADS15*) a member of a group of four rice genes that play similar roles in specifying inflorescence meristem identity (Kobayashi et al., 2012). *SbMADS7* is also co-orthologous to two maize genes *zap1* and *mads3*. *Zap1* was identified in a similar transcriptome-wide association study for flowering time variation in maize (Torres-Rodríguez et al., 2024). The second MADS-box gene identified, *SbMADS31*, appears to completely lack orthologs in maize (Figure 3A,B) but belongs to the clade of SOC1-like genes present in grasses. However, beyond members of the FT and MADS-box gene families, which have been linked to variation in flowering time in a wide range of species(Kobayashi et al., 1999, Meng et al., 2011, Corbesier et al., 2007, Becker and Theißen, 2003, Zhao et al., 2011), our analysis also linked multiple components of the aging pathway to variation in non-photoperiod sensitive sorghum.

The single gene whose expression was most significantly associated with flowering time in temperate sorghum was *SbSBP19* (Figure 2A). Loss of function alleles maize ortholog of this gene (*ZmSPL29*) have been shown to delay flowering in maize, with additional synergistic effects on flowering time in double mutants for *ZmSPL29* and *ZmSPL13* (Yang et al., 2023). *ZmSPL13* is ortholog of *SbSBP4*, another genes whose expression is strongly linked to flowering time in sorghum (Figure 2A). Many SBP transcription factors are targeted by mir156 (Chuck et al., 2007) and we computationally confirmed that both *SbSPB19* and *SbSPB4* are targets of mir156 in sorghum. Despite the lack of microRNA gene model annotations in the Sorghum v5 reference genome, we identified a strong signal from a gene model that, despite being annotated as a protein coding gene, corresponds to a copy of mir156 (Figure 2B) as well as a suggestive signal from a second gene model which also corresponds to a copy of mir156. The microRNA mir156 regulates vegetative phase change in maize and arabidopsis and overexpression of mir156 delays flowering in both species (Wu and Poethig, 2006, Chuck et al., 2007). This microRNA also serves as the first step of the canonical aging flowering time pathway, with expression of mir156 declining over time, resulting in increases in the expression of SBP transcription factors which, in turn, promote adult vegetative identity. In maize, *ZmSPL13* and *ZmSPL29* directly bind to the promoter of and activate the expression of mir172 (Yang et al., 2023). The genomic sequence encoding sbi-MIR172a is located 41 kilobases from the strong flowering time GWAS hit located on chromosome 9 (Figure 1A). Computational prediction of mir172 targets in sorghum identified a number of genes including Sobic.001G036800, an AP2 transcription factor orthologous to the major maize flowering time regulator *rap2.7* which acts as a negative regulator of flowering time (Salvi et al., 2007).

The expression of a number of additional genes were linked to variation in flowering time in one or both environments but could either function via indirect mechanisms, act on flowering time in ways not yet characterized or not yet well characterized, or represent false positive associations. The sorghum ortholog of the maize gene *teosinte branched1* which functions in the control of axillary meristem dormancy or activation in both maize and sorghum (Doebley et al., 1995, Kebrom et al., 2006) was also linked to flowering time variation via TWAS (in both environments) as was the sole sorghum ortholog of the arabidopsis gene *tornado1* which regulates cell proliferation and differentiation in the meristem and the early stages of differentiating organs (Cnops et al., 2006). The expression of *SbNAC32* in leaf tissue was significantly associated with flowering time. Loss of function alleles of maize ortholog of *SbNAC32*, *ZmNAC132*, exhibit delayed floral development (Yuan et al., 2023). However the gene also appears to play a key role in regulating leaf senescence (Yuan et al., 2023), suggesting a potential alternative model where, while the association between the expression of *SbNAC32* and flowering time does represent a causal relationship, variation in flowering time may be driving differences in senescence and hence gene expression, rather than vice versa. The three additional genomic intervals which were linked to variation in flowering time in a single environment via a multi-locus model but not supported in the second environment may represent loci which act on flowering time only in specific environmental conditions. Alternative it may be that the effect of these loci was overpowered by the three larger effect loci in the single locus testing model employed for second environment validation. Ultimately field experiments employing the entire larger sorghum population, which can support multi-locus GWAS approaches at each site, could be used to address this question.

Overall this study was able to identify a substantial number of strongly supported genes and genomic intervals associated with variation in flowering time in a largely photoperiod-insensitive diversity panel sorghum converted to flower in temperate environments. Several factors likely explain the success of this approach including the expanded size of the population employed which was approximately twice the size of the most widely used sorghum association panel (Casa et al., 2008). When we constrained our analysis to only those sorghum genotypes included in the sorghum association panel we were unable to identify any significant signals for flowering time (Figure S4), although this result should be treated with caution as we had genetic marker and trait data for only 303 sorghum association panel members, while typical past studies might have included approximately 350 members of the panel (Mural et al., 2021). The set of more than 170,000 markers present at minor allele frequencies greater than 0.05 and located primarily in exonic sequences generated as part of this study represents a significant increase in power to detect marker trait associations from the previous set of approximately 80,000 markers scored for sorghum conversion lines which included many extremely rare alleles (minor allele frequencies between 0.025-0.05) and many genetic markers located in intergenic regions (Griebel et al., 2021) as the average intergenic genetic marker is less likely to be associated with functional variants (Rodgers-Melnick et al., 2016). Similarly, while at least one previous transcriptome-wide association study has been conducted in sorghum (Ferguson et al., 2021), the earlier study employed RNA-seq from only 229 sorghum genotypes and utilized TWAS primarily to support moderate GWAS signals rather than as an independent gene discovery method. Here we generated data from more than three times as many sorghum genotypes, collected from a single field experiment in a time window of less than two hours, reducing confounding sources of transcriptional variation which can reduce power for detecting transcript-phenotype associations (Torres-Rodríguez et al., 2025), which make explain the large number of well supported trait-associated genes identified in this study. Several recent studies have demonstrated that gene expression data can be reused to discover genes controlling traits scored in other environments and expressed in other tissues (Li et al., 2021, 2024, Torres-Rodríguez et al., 2024). As a result, both genetic marker data and transcript abundance data generated for this larger population of sorghum conversion lines have the potential to enable and accelerate gene discovery for a wide range of traits and use cases.

## Acknowledgments

The authors thank Christine Smith and Brandi Sigmon for assistant with field design and maintenance in Nebraska. They thank Mackenzie Malcom nee Zwiener, Han Tran, Isabel Sigmon, Alliance Igiraneza, Aleah Miller, Lionel Kagaba, Babou Clion Muhoza, Mohamed Al Hussaini, Prince Ngiruwonsanga, Henry Medlock, Daniella Norah Tumusiine, Grace Carey, Annika Waller, Theresa Tran, Alice Guo, Alexis Finch for their contribution to the planting, weeding, and trait data collection of the Nebraska field site. The authors thank Fangyi Li, Marcin Grzybowski, Hongyu Jin, Ravi Mural, Guangchao Sun, Alliance Igiraneza, Michael Tross, Nathan Korth, Mackenzie Malcom nee Zwiener for their contribution to gene expression profiling. The authors also thank Jarius Whitehead for assistance with data collection in Alabama.

This work was supported by the U.S. Department of Energy, Grants no. DE-SC0020355 and DE-SC0023138, the National Science Foundation under grant OIA-1826781, USDA-NIFA under the AI Institute: for Resilient Agriculture, Award No. 2021-67021-35329 and the Foundation for Food and Agriculture Research Award No. 602757 to JCS.

## Author contributions statement

HM, JVTR, and JCS conceived of the experiments. KL, JT, BL, EC, and XK generated the data. HM and NS processed the data. HM and JVTR conducted the experiments and visualized the results with input from JCS and NS. HM and JCS wrote the initial draft of the manuscript. All authors reviewed and approved the final version of the manuscript.

## Data availability

RNA-seq data generated as part of this study has been deposited in the European Nucleotide Archive (ENA) under study accession number PRJEB83049. SNP calls and processed expression data have been uploaded to Figshare: **10.6084/m9.figshare.27936195**. Scripts and code used to conduct analyses and visualize data have been uploaded to the github repository associated with this study, as have plot level trait measurements and processed trait BLUEs. The complete set of GWAS hits identified in Nebraska and the complete set of TWAS results, including SNPs and genes which did not exceed the cutoff for significance have been uploaded to the github repository associated with this paper.

## Additional Information

### Supplementary Figures

**Figure S1.**
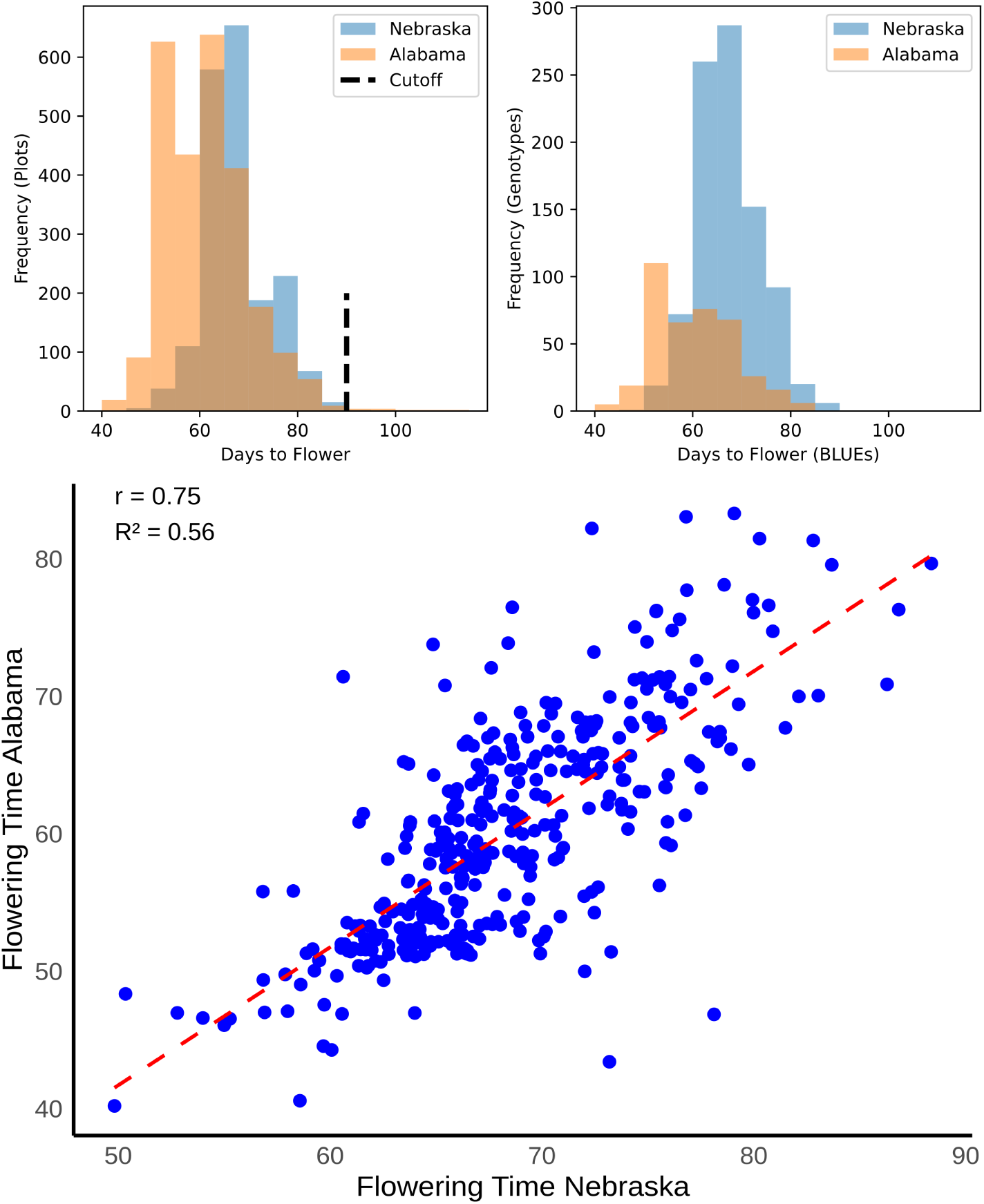
Distributions and correlations flowering time within and between Nebraska and Alabama. Top left shows the distribution of non-spatially corrected flowering times recorded for each plot in each environment. Vertical dashed black line indicates the cutoff used to exclude extremely late flowering plots. Top right shows the distribution of spatially corrected BLUEs for each genotype in each environment. Bottom shows the relationship between spatially corrected best unbiased linear estimators in the Alabama (y-axis) and Nebraska (x-axis) field experiments. A dashed red line indicates the best fit linear regression. N = 365 (for the genotypes shared between the Nebraska and Alabama)

**Figure S2.**
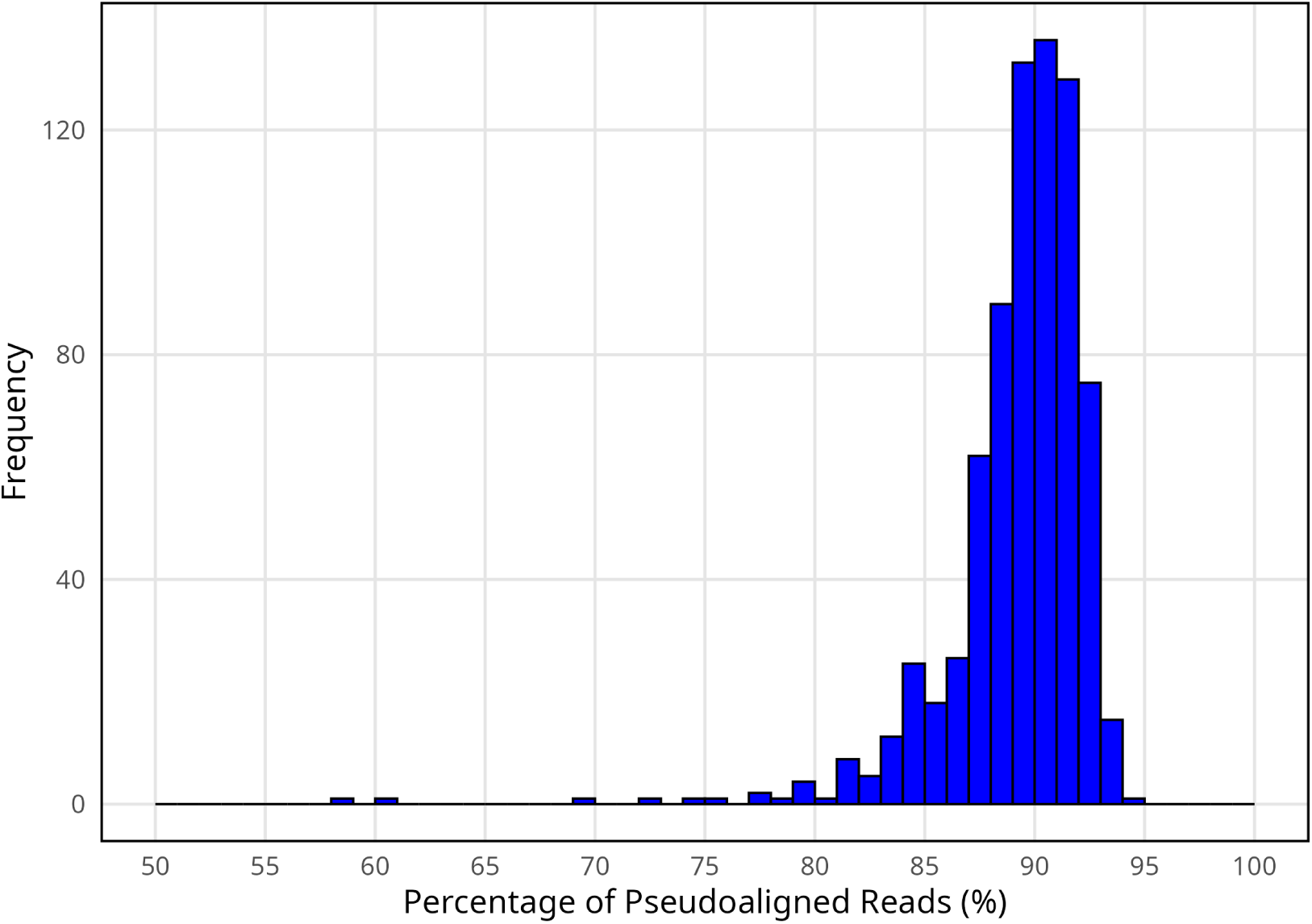
Distribution of pseudoalignment rates for each RNA-seq sample used in this study. Proportion of reads that could be pseudoaligned to the “primary transcript only” transcript sequences of the BTx623 v5 sorghum reference genome via kallisto. One extremely low alignment sample, PI 656042, where only 8.9% of reads could be matched to sorghum transcripts, is omitted from this visualization to aid in readability.

**Figure S3.**
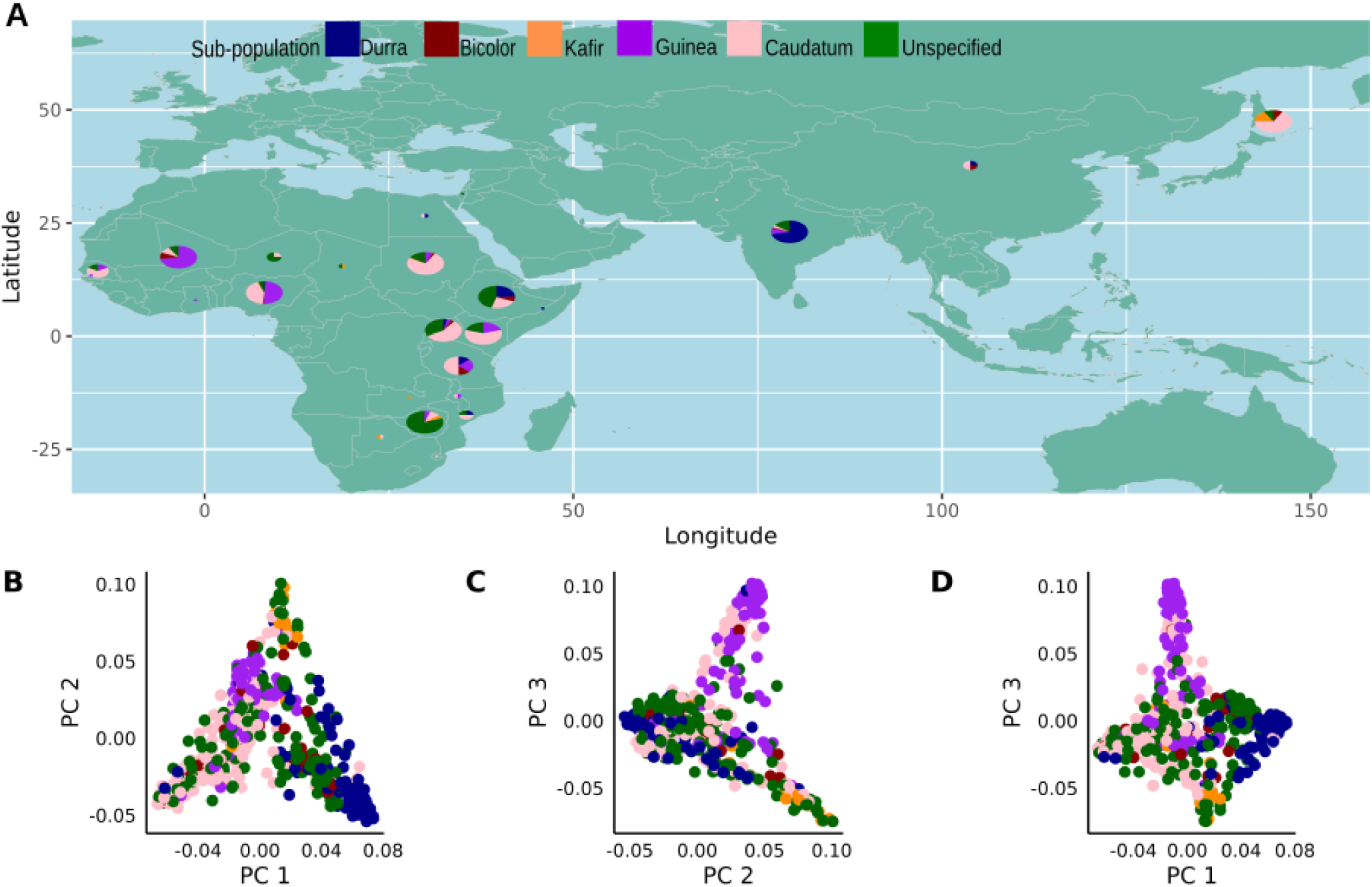
Sorghum around the world and PCs with relation to the sub-population of sorghum: A. World Map for the Sorghum Diversity Panel and the places around the world with respect to the sub-population. The Principal Components are generated using SNP dataset B. Principal Component 1 with respect to Principal Component 2 with the colors defining sub-population. C. Principal Component 2 with respect to Principal Component 3 with the colors defining sub-population. D. Principal Component 3 with respect to Principal Component 1 with the colors defining sub-population.

**Figure S4.**
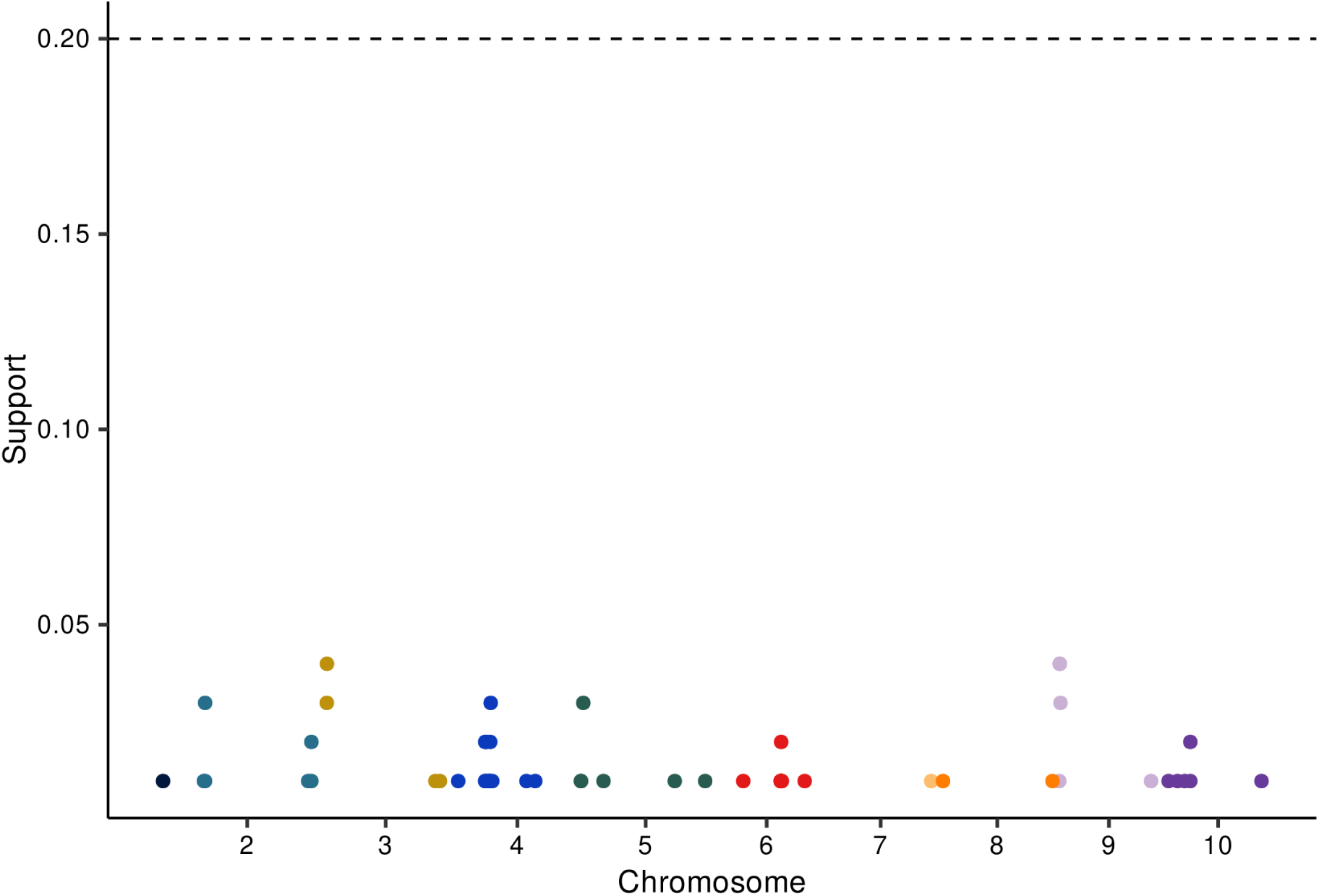
Results of a genome wide association study conducted using only lines from the Sorghum Association Panel. Genome wide association study conducted for flowering time measured across 303 sorghum genotypes scored in Lincoln, Nebraska in 2021 which are also part of the Sorghum Association Panel. Y-axis indicates the resampling model inclusion probability calculated from 100 iterations of the FarmCPU GWAS algorithm. Horizontal dashed line indicates the threshold employed to consider a marker significantly associated with flowering time in this study (RMIP = 0.2).

**Figure S5.**
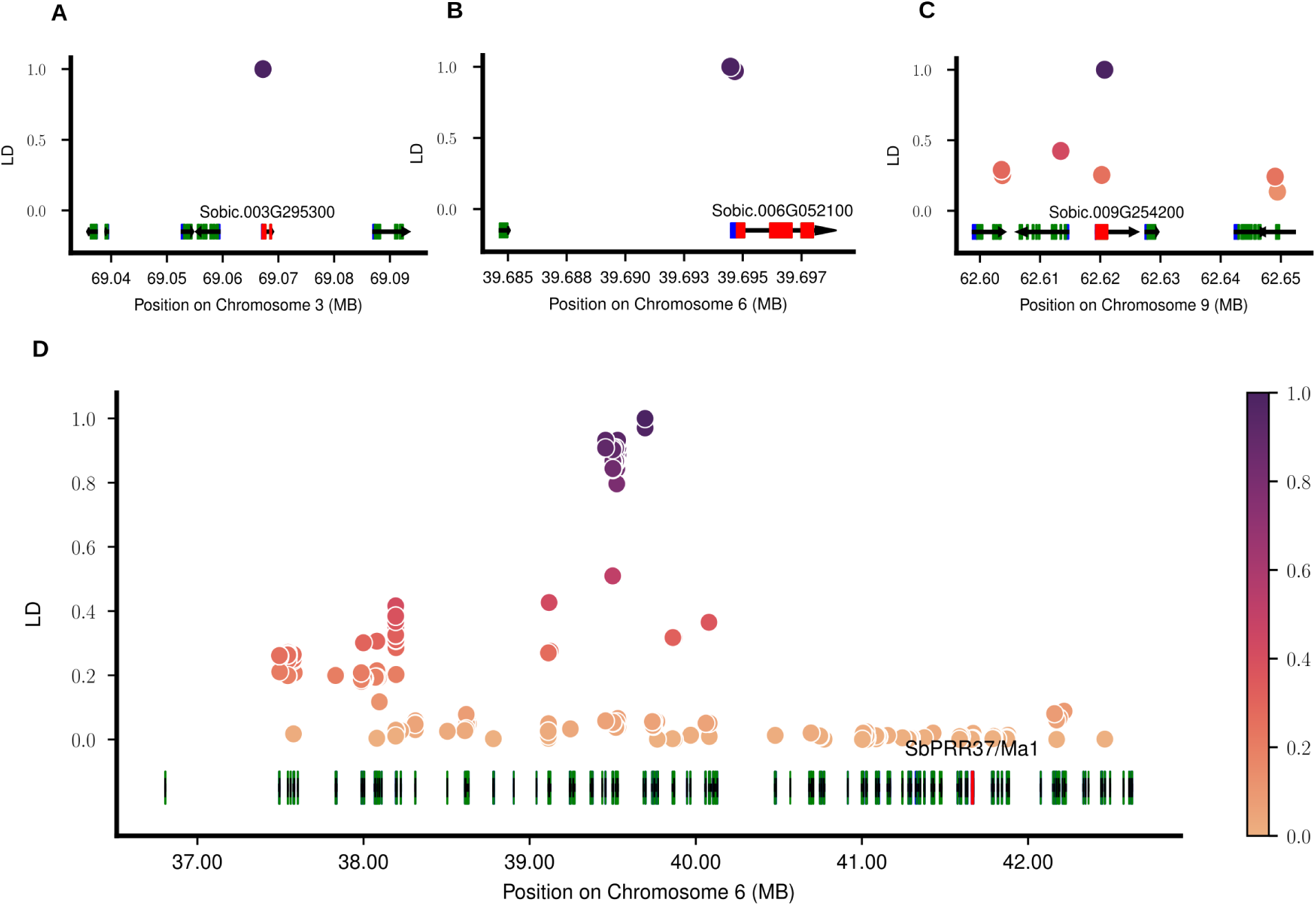
Linkage Disequilibrium for the most significant SNPs found in the GWAS analysis. **A)** Genomic interval and annotated genes surrounding the trait associated marker Chr03:69,067,236. Gene containing the marker indicated in red, other genes within the interval shown in green (protein-coding exons) and blue (untranslated exons). Each genetic marker within the interval is indicated with a circle whose color and position on the y-axis correspond to the degree of linkage disequilibrium between that marker and the trait associated marker. **B)** Genomic interval and annotated genes surrounding the trait associated marker Chr06:39,694,475. **C)** Genomic interval and annotated genes surrounding the trait associated marker Chr09:62,620,720. **D)** An expanded view of the six megabase genomic interval surrounding the flowering time associated marker Chr06:39,694,475. The position of Sobic.006G057866 the gene model corresponding to SbPRR37 thought to correspond to *Maturity1* is indicated in red. Exons of genes within the interval are shown in green (protein coding exons) and blue (untranslated exons). Each genetic marker within the interval is indicated with a circle whose color and position on the y-axis corresponds to the degree of linkage disequilibrium between that marker and Chr06:39,694,475.

**Figure S6.**
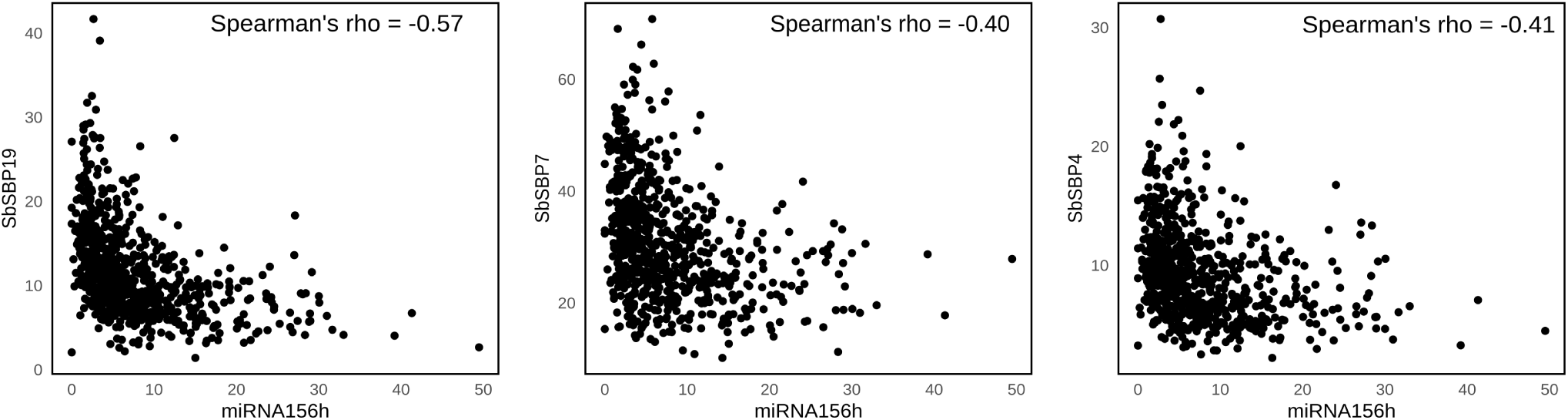
Correlation between the expression of sbi-MIR156h and the three SBP transcription factors identified via TWAS. Each point indicates the expression of sbi-MIR156h in TPM (x-axis) and the expression of the SBP transcription factor name given on the y-axis in units of TPM in one of the 738 sorghum RNA-seq samples analyzed as part of this study.

**Figure S7.**
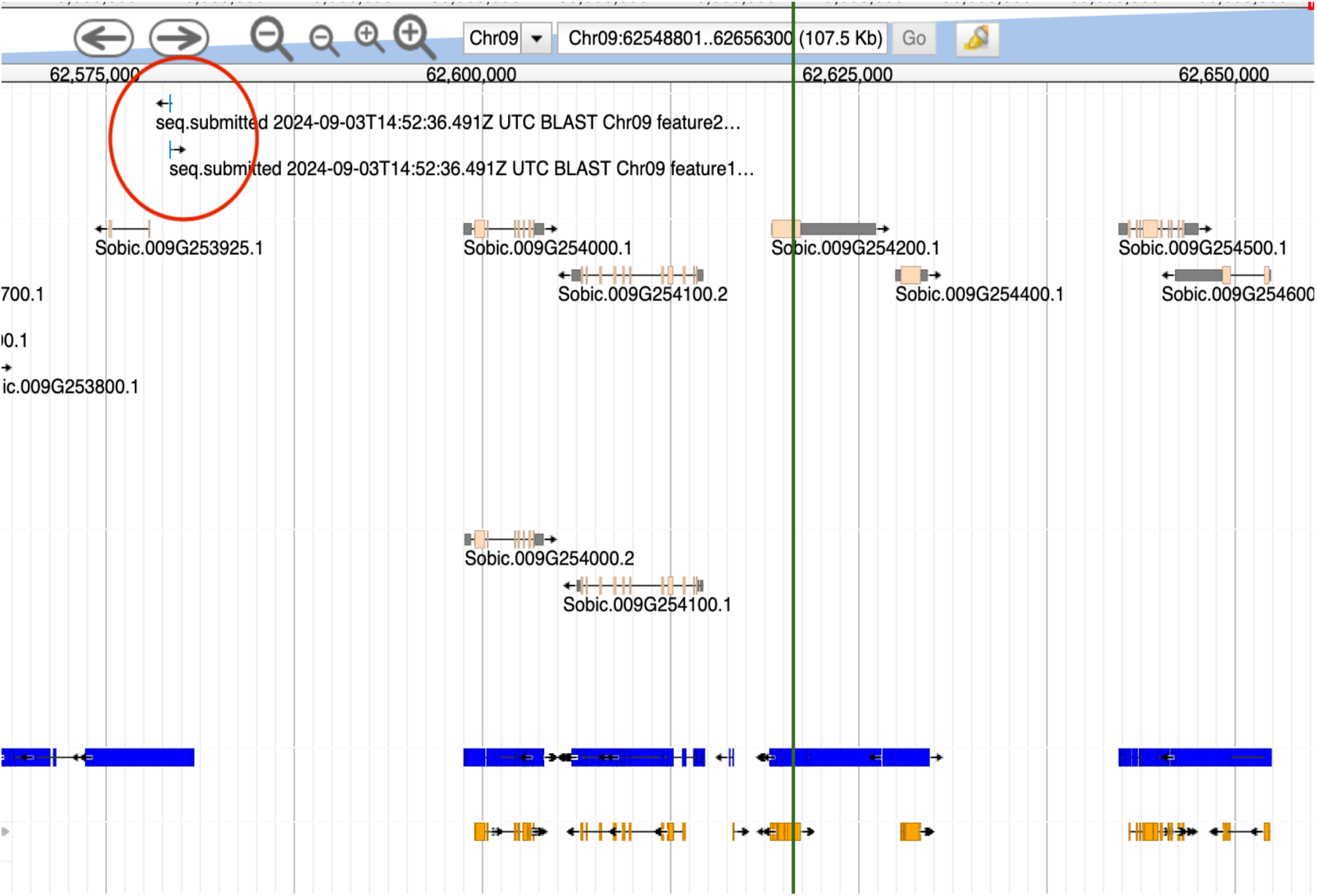
Position of mir172a relative to the flowering time associated marker Chr09:62,620,720. Genomic interval and annotated genes surrounding the trait associated marker Chr09:62,620,720. Screenshot from the phytozome genome browser showing the position of mir172a (red circle) and the position of Chr09:62,620,72 green verticle line.

**Figure S8.**
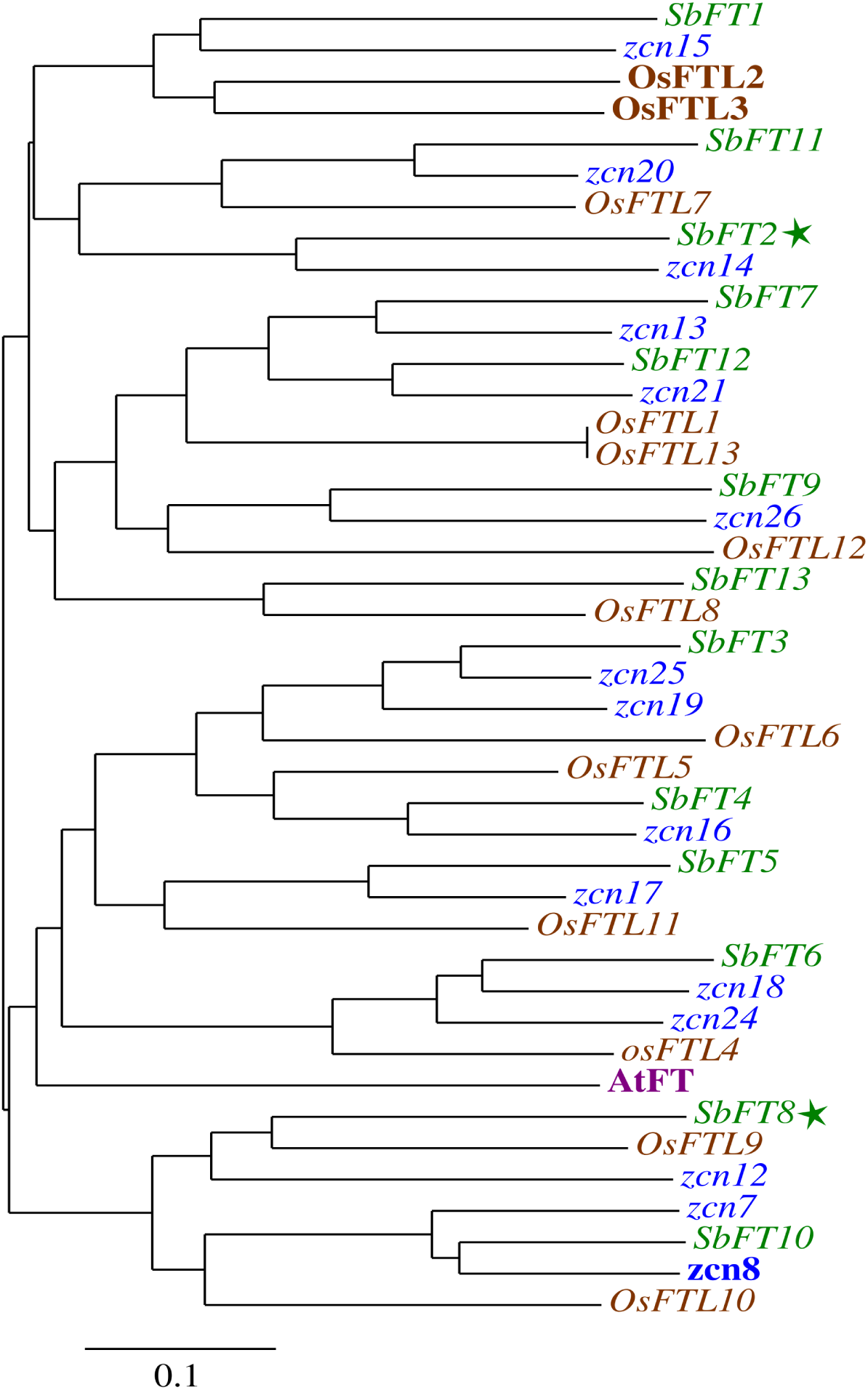
Phylogenetic tree for FT genes. Phyogenetic relationships among the set of FT-like genes found in the sorghum, maize, rice, and arabidopsis genomes. Star indicates the sorghum gene copy linked to variation in flowering time via TWAS and GWAS in our study. Phylogeny constructed using PhyML using a multiple sequence generated by aligning the nucleotide sequence of the primary transcript for each gene with MUSCLE. Bold names indicate the direct promoter of flowering-time in the specific

### Supplementary Tables

**Supplemental Table S1.** Annotated sorghum gene models and functional annotations located within 100 kilobases of the flowering time associated genetic marker located at Chr09:62,620,720.

| GeneID | Function |
| --- | --- |
| Sobic.009G253000 | RNA polymerase sigma factor |
| Sobic.009G253101 | Pentatricopeptide repeat containing protein |
| Sobic.009G253150 | Pentatricopeptide repeat containing protein |
| Sobic.009G253200 | RNA polymerase sigma factor |
| Sobic.009G253300 | MYB transcription factor |
| Sobic.009G253400 | Hyaluronan (RNA binding) |
| Sobic.009G253450 | Unknown protein, no homologs |
| Sobic.009G253500 | F-box domain containing protein |
| Sobic.009G253600 | Pentatricopeptide repeat containing protein |
| Sobic.009G253700 | Phosphatidylinositol 4-kinase |
| Sobic.009G253800 | Aspartyl protease |
| Sobic.009G253925 | Unknown protein, no homologs |
| Sobic.009G254000 | Protein kinase |
| Sobic.009G254100 | RNA helicase |
| Sobic.009G254200 | Protein kinase |
| Sobic.009G254400 | Leucine rich repeat protein. |
| Sobic.009G254500 | Protein kinase |
| Sobic.009G254600 | Alpha/beta hydrolase |
| Sobic.009G254700 | Alpha/beta hydrolase fold-containing protein |
| Sobic.009G254750 | Unknown protein, no homologs |
| Sobic.009G254800 | Heat shock protein |
| Sobic.009G254900 | Lysine decarboxylase |
| Sobic.009G255000 | Protein kinase |
| Sobic.009G255100 | Membrane protein homologous to hypersensitive response induced gene |

**Supplemental Table S2.** Names and gene model identifiers for the thirteen FT-like sorghum genes described by (Wolabu et al., 2016), as well as names and gene model identifiers for syntenic orthologs of these sorghum genes in the genomes of rice, and both maize subgenomes.

| Sorghum (v5) | Rice | Maize1 | Maize2 | Sorghum (v1) |
| --- | --- | --- | --- | --- |
| SbFT1<br>Sobic.010G045100 | osFTL2/Hd3a<br>LOC_Os06g06320<br>osFTL3<br>LOC_Os06g06300 | NA | zcn15<br>Zm00001eb271180 | Sb10g003940 |
| SbFT2<br>Sobic.003G017200 | osFTL1<br>LOC_Os01g11940 | NA | zcn14<br>Zm00001eb338650 | Sb03g001700 |
| SbFT3<br>Sobic.006G128500 | osFTL6<br>LOC_Os04g41130 | zcn25<br>Zm00001eb078980 | zcn19<br>Zm00001eb425430 | Sb06g020850 |
| SbFT4<br>Sobic.004G206600 | osFTL5<br>LOC_Os02g39064 | zcn16<br>Zm00001eb246470 | NA | Sb04g025210 |
| SbFT5<br>Sobic.005G110406 | osFTL11<br>LOC_Os11g18870 | zcn17<br>Zm00001eb090320 | NA | Sb0010s003120 |
| SbFT6<br>Sobic.002G262500 | osFTL4<br>LOC_Os09g33850 | zcn24<br>Zm00001eb318290 | zcn18<br>Zm00001eb102650 | Sb02g029725 |
| SbFT7<br>Sobic.004G101800 | osFTL13<br>LOC_Os02g13830 | zcn13<br>Zm00001eb239360 | NA | Sb04g008320 |
| SbFT8<br>Sobic.003G295300 | osFTL9<br>LOC_Os01g54490 | zcn12<br>Zm00001eb153190 | NA | Sb03g034580 |
| SbFT9<br>Sobic.010G164200 | osFTL12<br>LOC_Os06g35940 | zcn26<br>Zm00001eb384770 | NA | Sb10g021790 |
| SbFT10<br>Sobic.009G199900 | osFTL10<br>LOC_Os05g44180 | zcn7<br>Zm00001eb293080 | zcn8<br>Zm00001eb353250 | Sb09g025760 |
| SbFT11<br>Sobic.008G082200 | osFTL7<br>LOC_Os12g13030 | zcn20<br>Zm00001eb411580 | NA | Sb08g008180 |
| SbFT12<br>Sobic.006G047700 | osFTL13<br>LOC_Os02g13830 | zcn21<br>Zm00001eb085740 | NA | Sb06g012260 |
| SbFT13<br>Sobic.003G026600 | osFTL8<br>LOC_Os01g10590 | NA | NA | Sb03g002500 |

**Supplemental Table S3.**
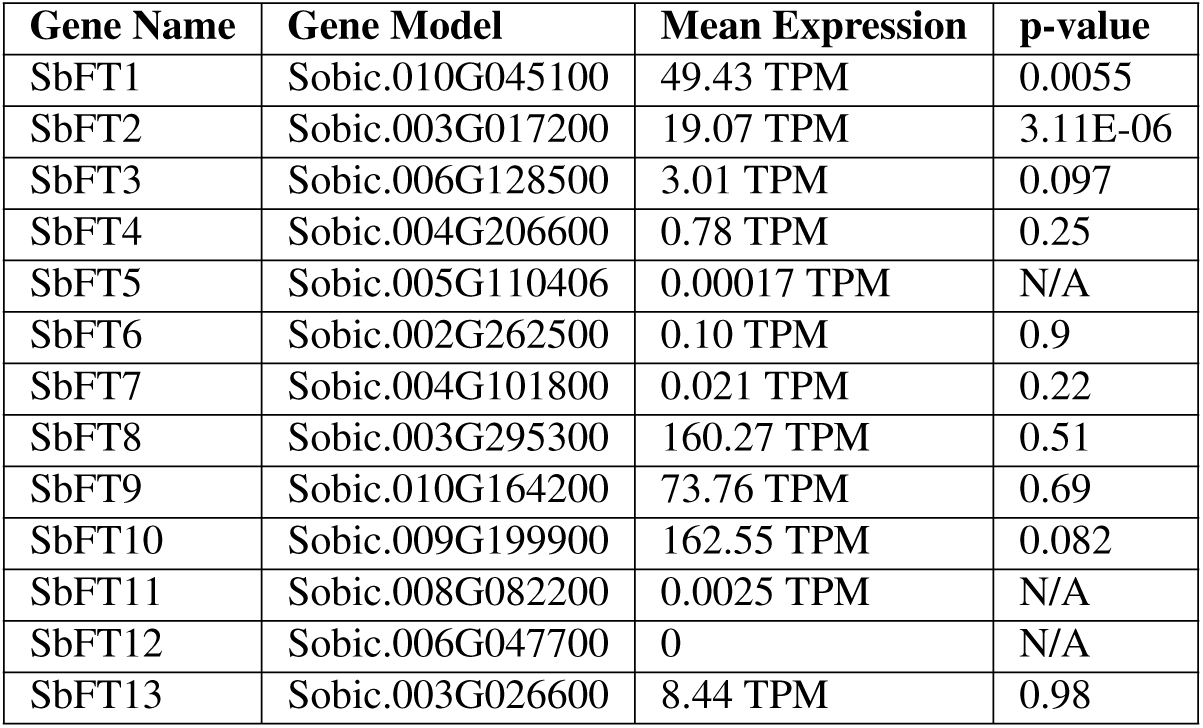
Average expression level and significance of association with flowering time for all annotated FT-like genes in the sorghum genome.

